# ProDive: detecting cross-family local conservation

**DOI:** 10.64898/2026.04.02.715841

**Authors:** Xiang Chen, Pu Tian

## Abstract

Short conserved sequence–structure elements can recur across protein families even when whole-domain homology is weak or absent, but systematic detection of such short local signals remains difficult. Here we present ProDive, a profile hidden Markov model (pHMM)-based framework for detecting cross-family local conservation. ProDive combines a closed-form approximation to the divergence between equal-length pHMM windows, a background-suppression stage that identifies and removes low-information similarities, diagonal-path extraction in window-pair similarity landscapes, and filtering and rescoring of retained paths according to the fraction of seed-alignment sequences supporting the matched positions. GPU benchmarking showed numerical agreement with CPU calculations and predictable near-linear scaling with the number of window-pair matrix elements. When applied to all 25,545 Pfam families in release 38.0, ProDive yielded 318,289 high-confidence pHMM segment correspondences after filtering. For fragment pairs with suitable representative structures, length-stratified controls showed that ProDive matches in the 8–13-aligned-residue regime have significantly lower Cα root-mean-square deviation (RMSD) than same-length random fragment pairs. Comparison with HHsearch showed that ProDive occupies a distinct fragment-scale detection niche: ProDive paths were largely outside HHsearch-reported family-pair space, and in shared family pairs ProDive boundaries isolated shorter compact cores. Together, these results establish ProDive as a practical method for local conservation identification and a resource for downstream studies of protein structure, evolution, and design.

## 1 Introduction

Protein similarity is typically analyzed at the level of domains and folds. Foundational resources such as SCOP, CATH, Pfam, and ECOD capture these relationships at multiple structural and evolutionary scales. ^1–4^ Structure-comparison tools such as DALI, TM-align, and Foldseek, together with profile– profile methods such as HHsearch, have made remote homology detection and structure-guided annotation highly effective for family- and fold-level problems. ^5–8^ In contrast, much less methodological attention has been devoted to the detection of short local conservation signals that recur across otherwise distinct protein families. Several studies have argued that proteins reuse short building blocks, or fragment vocabularies, and a number of local comparison strategies have been explored. ^9–16^ However, standard local sequence-alignment methods are not designed to compare two pHMM segments directly, and whole-profile methods tend either to miss weak local similarities or to merge them into longer, less specific pHMM segments.

The central technical obstacle has two parts. First, short cross-family signals are often too weak and degenerate to identify reliably from individual sequence–sequence comparisons alone. Representing each family as a pHMM helps by pooling family-level variation and retaining statistical evidence for local patterns. However, a pHMM defines a probability distribution over an effectively infinite sequence space, so exact sequence-distribution divergence would require summing over an intractable set of emitted sequences and paths. A common tractable substitute in profile–profile comparison is to optimize and report a single high-probability profile-alignment path. This path-based summary is powerful for remote-homology searches such as HHsearch, where the goal is to identify a coherent profile-level relationship, often over a longer interval. For short local correspondence signals embedded in diffuse family distributions, however, a single optimized path can either miss the signal or absorb it into a broader alignment that does not isolate the conserved local segment. A practical fragment-scale comparison therefore requires a tractable approximation that incorporates the probability distributions encoded by both profiles without reducing their comparison to a single optimal path. Here we introduce ProDive (Profile HMM Divergence), which addresses this problem through a closed-form approximation to the divergence between equal-length pHMM windows. Sliding windows extend the comparison to local segments of unequal-length profiles, while parallel evaluation, background suppression, diagonal-path extraction, and seed-alignment-support filtering and rescoring make database-scale segment detection practical.

Throughout, segment correspondence refers to similarity defined at the pHMM level, whereas fragment pair refers to its representative-chain realization after projection through seed alignments. A window is a fixed-length span of *k* pHMM state positions, and a diagonal path is a stretch of contiguous high-scoring window pairs defining matched pHMM segments. Structural characterization is post-detection and is described in Methods; aligned lengths and RMSD values refer only to structurally characterized representative-chain fragments. We apply ProDive to all Pfam families and assess the resulting pHMM segment correspondences through structural validation and comparison with HHsearch. The analysis yields 318,289 high-confidence correspondences, with structural validation supporting the detected local similarities and reciprocal comparison with HHsearch showing asymmetric but limited overlap: most ProDive paths occurred outside HHsearch-reported family-pair space, and most HHsearch-reported family pairs lacked an overlapping ProDive path.

## 2 Results

### 2.1 Derivation of the ProDive formulation and its application to Pfam families

The central algorithmic contribution of ProDive is a closed-form approximation to the full probabilistic divergence between two equal-length pHMMs. Starting from the HMM divergence heuristic proposed by Mohammad and Tranter,^17^ and using Markov-source KL-divergence-rate theory as a related reference point, ^18^ we exploit the structure of the pHMM transition matrix to derive an asymptotic expression for the directional divergence between two models with parameter sets *λ* and *λ ′*:

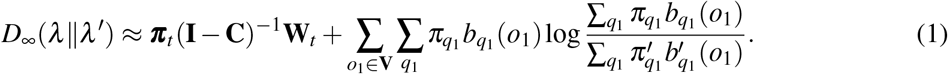

Here ***π***_*t*_ is the initial transient-state distribution, **C** is the principal submatrix, and **W**_*t*_ collects statespecific emission-divergence terms. The full explanation of the notation and complete derivation is provided in the Supplementary Notation and Supplementary Methods Section S1. To compare pHMMs of unequal lengths, each model is partitioned into overlapping windows of length *k*. ProDive uses the average of the two directions to remove divergence asymmetry, and normalizes by the number of basic terms accumulated over windows to yield a length-invariant score *D*_norm_ (Methods).

We applied this workflow to all 25,545 families of Pfam release 38.0 using seed alignments as the underlying family representation. ^3^ Evaluating *D*_norm_ across all window pairs produced a dense similarity landscape in which local similarity appeared as faint diagonal chains embedded in extensive background signal, as illustrated for the representative Pfam pair PF00593 and PF10677 (Fig. 1A). Initial candidate screening retained a subset of window pairs (Fig. 1B). Each candidate was then rescored by combining its pairwise divergence with the information content of the corresponding windows relative to the pHMM background distribution. This information-weighted landscape made broad low-information stripes explicit (Fig. 1C), whereas background suppression isolated sparse diagonal chains of local similarity (Fig. 1D) and produced a score landscape with a long, sparse right tail. The global distribution of this score showed that most window pairs remained near background values, whereas a small high-score tail emerged (Fig. 2A). Consecutive high-scoring points were linked into diagonal paths, each defining a cross-family segment correspondence between the two pHMMs. Because matched pHMM regions can differ in seed-alignment support, clean paths were further reweighted by the mean support of the two matched segments. This seed-support rescoring compressed weak paths before final filtering and defined the high-confidence correspondence set (Fig. 2B; Methods). After path-level quality filtering, the Pfam-wide scan yielded 318,289 high-confidence cross-family segment correspondences, which formed the input set for the structural validation and HHsearch comparison below.

**Figure 1.**
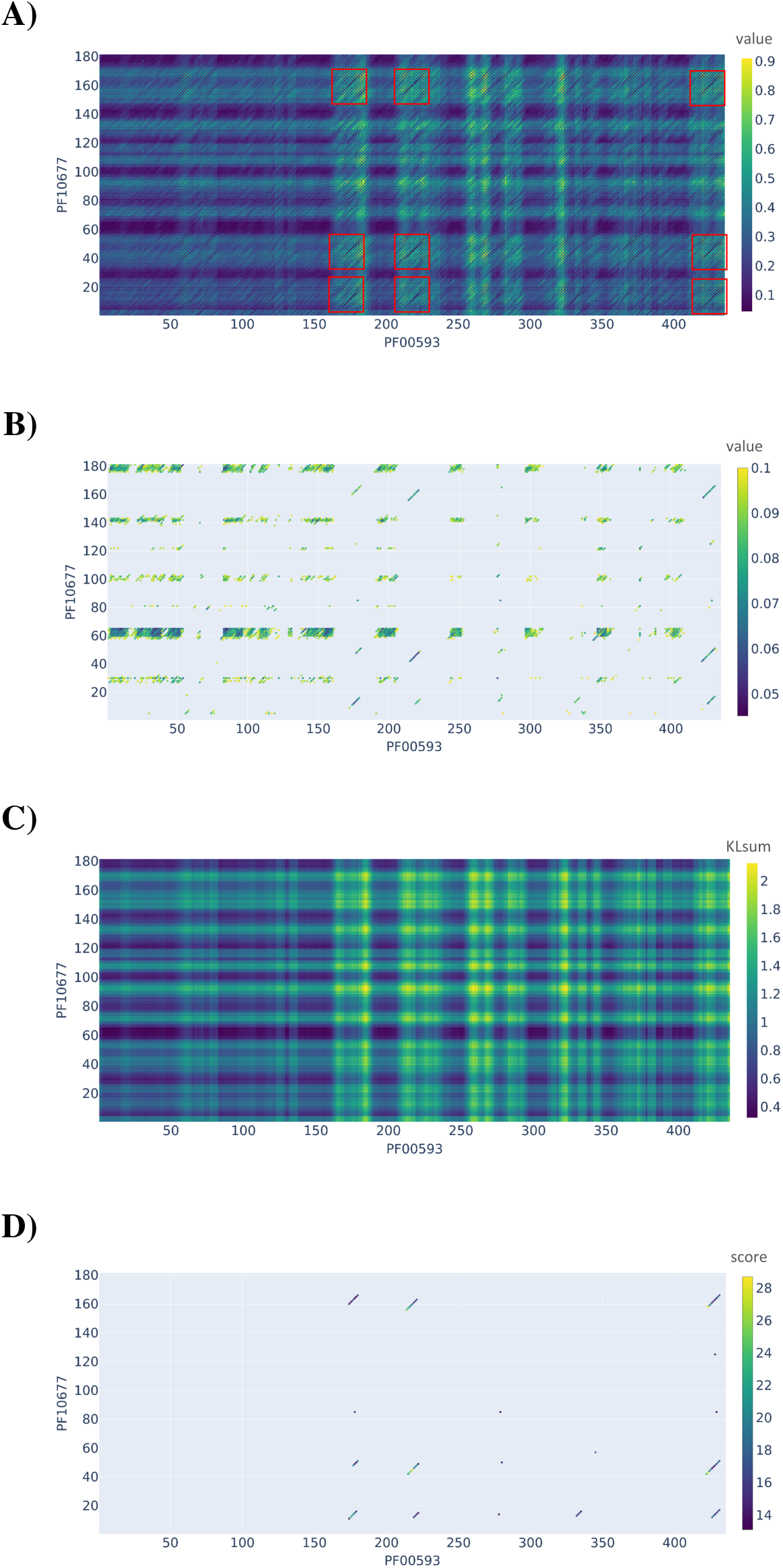
Representative four-stage visualization of a family-pair comparison from the Pfam-wide Pro-Dive scan, shown for PF00593 and PF10677 at window length *k* = 6. (A) Raw window-pair similarity landscape based on the symmetric divergence measure, with colorbar labeled `value`. (B) Candidate window pairs retained after the initial thresholding step (*D*_norm_ 0.1), again shown as `value`. (C) Background-information landscape across all window pairs, labeled `KLsum` and highlighting broad stripe-like patterns associated with nonspecific similarity. (D) Window pairs retained after background suppression and score filtering, labeled `score;` retained points above score 13 form sparse diagonal chains that define local segment correspondences.

**Figure 2.**
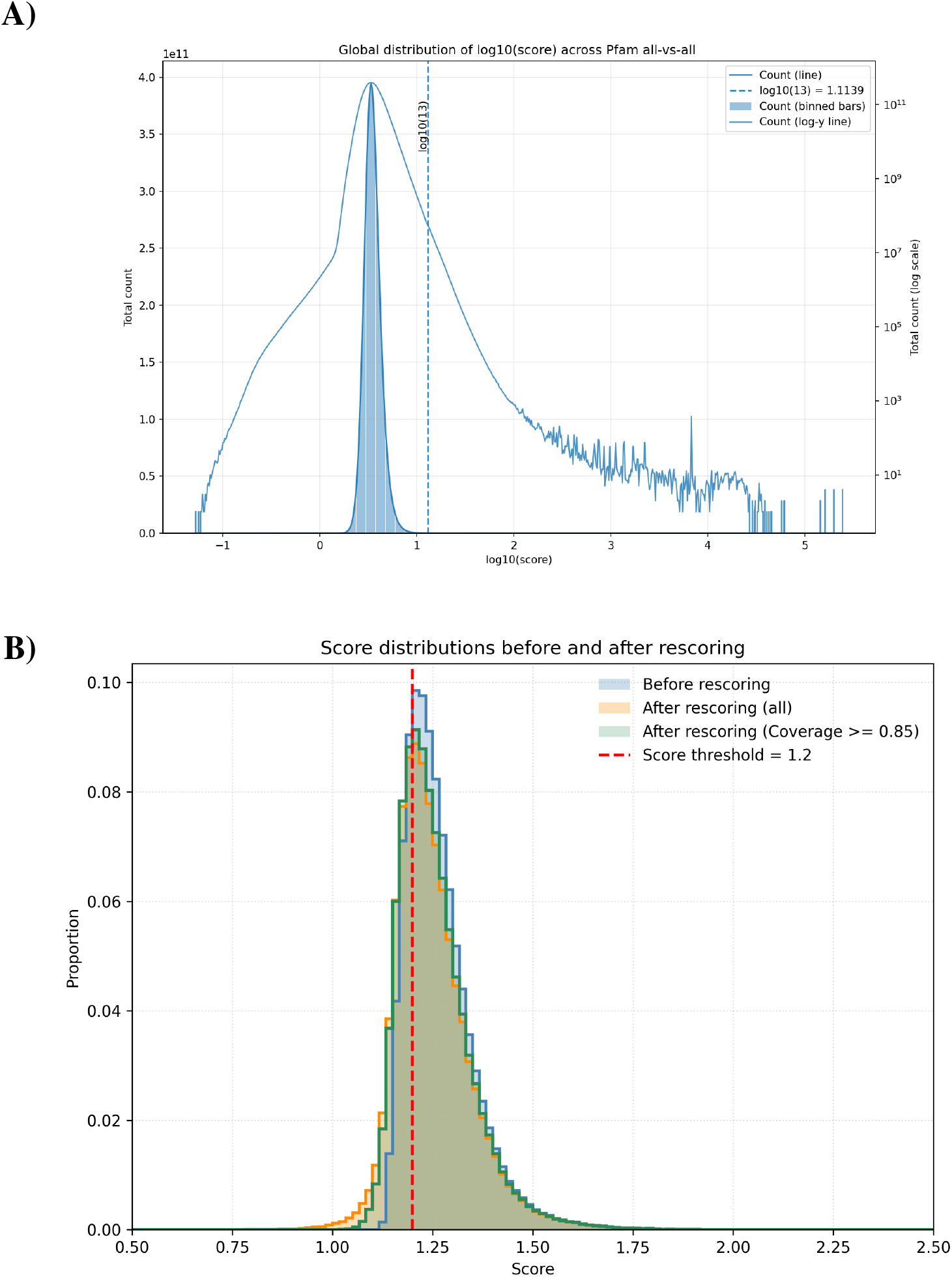
Filtering and rescoring behaviour in the Pfam-wide ProDive scan. (A) Global distribution of background-suppressed scores for individual window-pair candidate points, showing the sparse right tail above the operational point-score cutoff of 13. (B) Path-level score distributions before and after seedalignment-support rescoring, showing the effect of support reweighting and the final high-confidence path cutoff *S*_adj_ *>* 1.2.

### 2.2 Structural characterization and comparison with HHsearch

To assess whether retained ProDive paths correspond to local structural similarity rather than statistical noise, we projected pHMM segment correspondences onto representative sequences and calculated C*α* RMSD for mappable fragment pairs (Methods). We then compared ProDive and HHsearch by grouping Pfam family pairs as ProDive-only, shared, or HHsearch-only (Methods).

At the 20% HHsearch threshold, many ProDive-defined 8–13-residue fragment pairs mapped to family pairs not reported by HHsearch. Within this ProDive-unique class, representative-chain fragment pairs nevertheless showed low C*α* RMSD, indicating that the local ProDive correspondences remained structurally compact even when the corresponding family pair was absent from HHsearch results (Fig. 3A). Length-stratified random controls confirmed that this enrichment was not explained by chance structural similarity: across all tested core lengths from 8 to 13 residues, RMSD distributions of these ProDive fragment pairs were far below those of matched random fragment pairs, with *p*-values below machine precision (Supplementary Fig. S1). For family pairs reported by both methods, the structural distributions isolated the effect of segment-boundary definition because the family-level relationship was held fixed. ProDive-defined segments remained concentrated in the compact low-RMSD regime (Fig. 3B), whereas HHsearch-defined segments from the same family-pair class shifted toward longer alignments with broader RMSD values (Fig. 3C). Thus, ProDive-defined boundaries localized compact low-RMSD cores, whereas HHsearch-defined boundaries more often extended local similarity into longer and structurally broader intervals. Fragments projected from HHsearch-defined segments in family pairs without a ProDive path displayed the broadest and longest distribution (Fig. 3D); full-range views of the HHsearch-defined structural distributions are provided in Supplementary Fig. S2. The same distributional contrast persisted at the 70% HHsearch threshold (Supplementary Fig. S3A–C).

**Figure 3.**
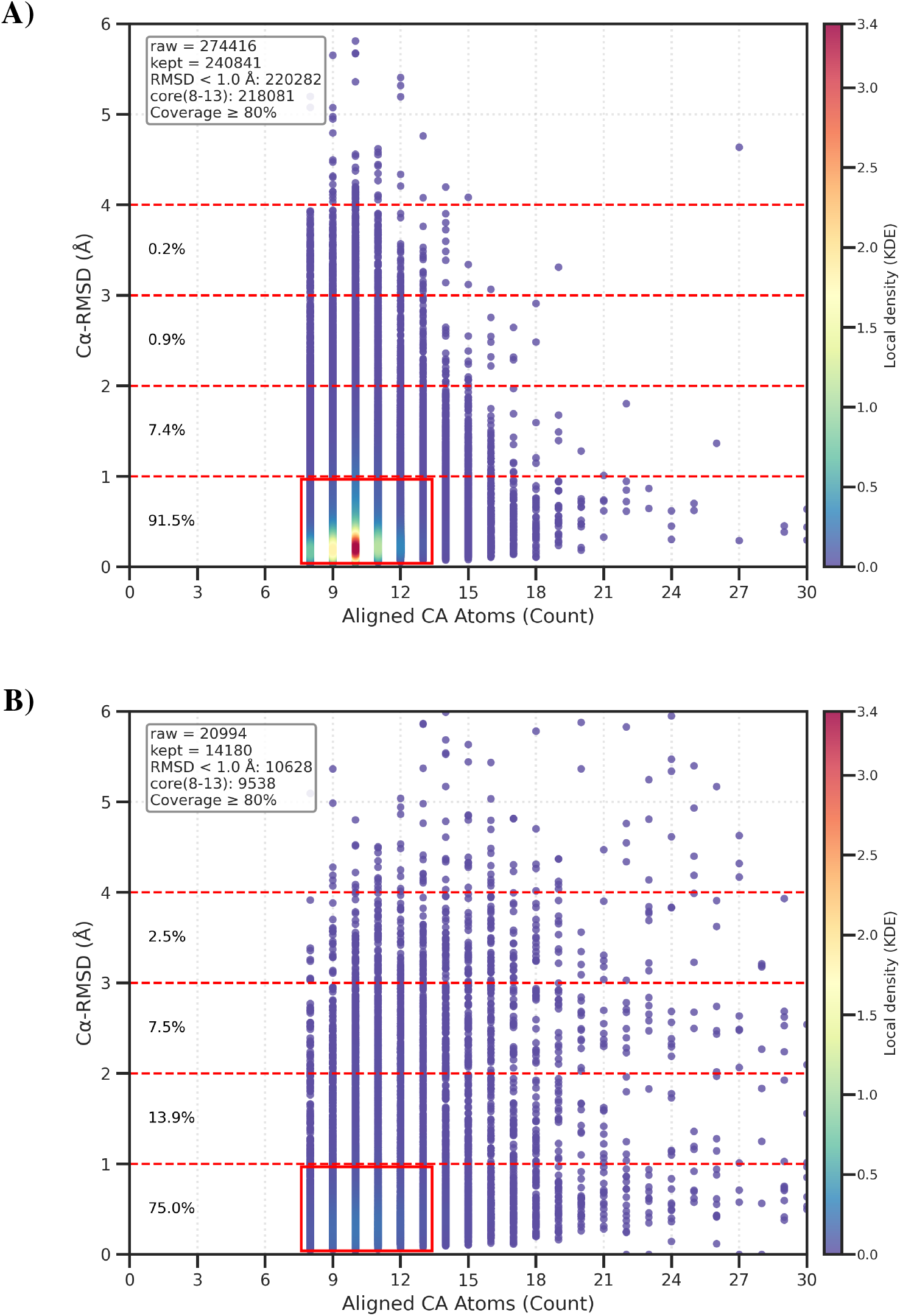
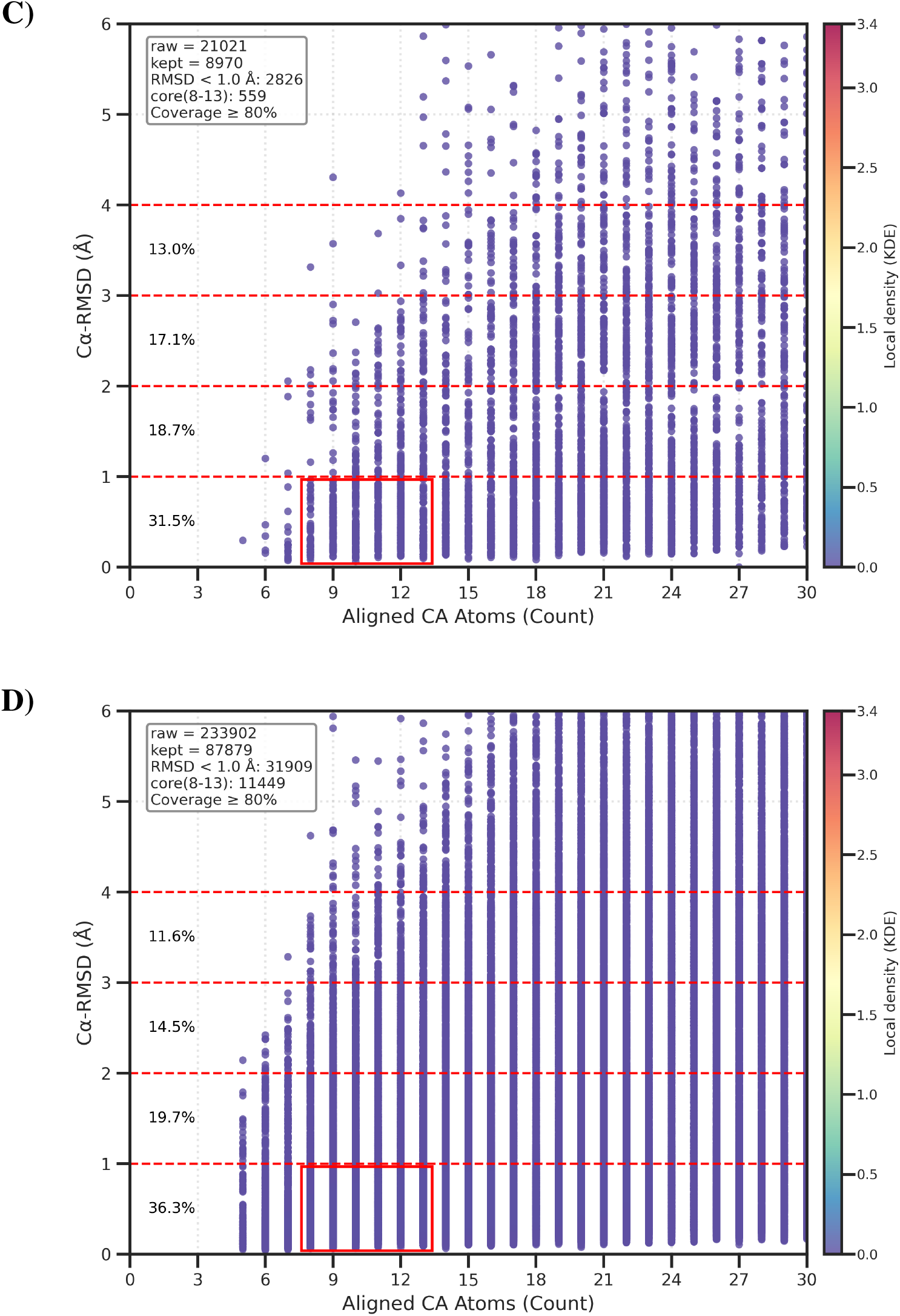
Post-detection structural characterization after classifying Pfam family pairs by HHsearch at the 20% probability threshold. Each panel shows representative-chain fragment pairs projected from method-defined pHMM segments as aligned C*α* atoms versus C*α* RMSD. The red box marks the compact local-core regime defined by 8–13 aligned C*α* atoms and RMSD < 1.0 Å. (A) ProDive-defined segments in ProDive-unique family pairs. (B) Shared family pairs evaluated with ProDive-defined boundaries. (C) Shared family pairs evaluated with HHsearch-defined boundaries. (D) HHsearch-defined segments in HHsearch-only family pairs.

This boundary contrast should be interpreted alongside a fundamental methodological distinction. HHsearch summarizes profile–profile comparison through an optimized high-probability alignment path, a path-based reduction well suited to detecting longer remote-homology relationships. ProDive was designed for a different regime: it scores fixed-length pHMM windows by an approximate divergence over the probability distributions encoded by both profiles, and only then links neighboring high-scoring windows into local paths. Thus, the HHsearch comparison is not simply a comparison of thresholds or boundary choices, but a comparison between path-based profile-alignment search and approximate full-profile local probability scoring. The present analysis does not prove that this algorithmic distinction alone causes the observed boundary distributions; it shows the practical consequence that the two methods occupy non-overlapping detection regimes.

When ProDive paths were stratified by ProDive-score cutoff, the ProDive-unique component was already present among the highest-scoring paths and increased as the cutoff was relaxed. At the 20% HHsearch threshold, paths from family pairs not reported by HHsearch accounted for 9.0k of 15.9k paths in the top 5% subset and 296.2k of 318.3k paths in the full filtered set; at the 70% threshold, the corresponding counts were 11.1k of 15.9k and 306.5k of 318.3k (Fig. 4A,B). A reciprocal analysis, using HHsearch-reported unique family pairs as the denominator, showed that the overlap remained a minority from the HHsearch perspective. At the 20% HHsearch threshold, 6.1k of 274.9k HHsearch family pairs overlapped the top 5% ProDive subset and 19.1k of 275.0k overlapped the full ProDive set; at the 70% threshold, the corresponding counts were 4.0k of 57.3k and 9.2k of 57.3k (Fig. 4C,D). Together, the random controls, structural comparisons, and reciprocal overlap analysis provide structural validation of ProDive at the dataset level and show that ProDive defines an independent fragment-scale detection regime distinct from domain-scale remote-homology search.

**Figure 4.**
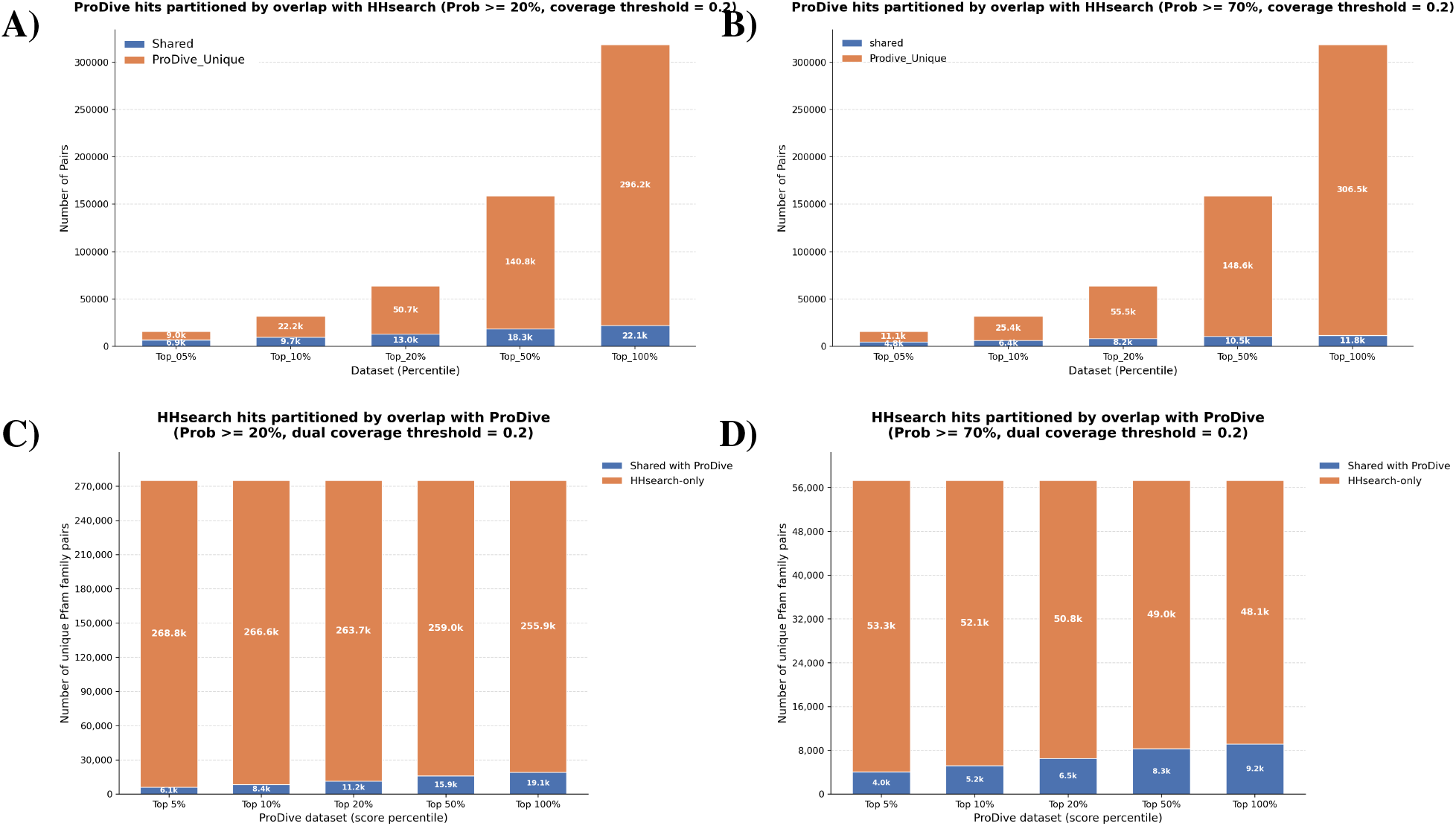
Reciprocal overlap between ProDive and HHsearch across ProDive-score cutoffs. (A,B) Pro-Dive paths were grouped by ProDive-score percentile cutoff (top 5%, 10%, 20%, 50%, and full set) and partitioned by whether their Pfam family pairs were reported by HHsearch at probability *≥* 20% (A) or *≥* 70% (B). (C,D) Unique HHsearch-reported Pfam family pairs at probability *≥* 20% (C) or *≥* 70% (C)were used as the denominator and partitioned by whether they overlapped a ProDive path from the indicated ProDive-score subset under a dual segment-coverage threshold of 0.2. The four reciprocal denominator-threshold combinations show that ProDive local-segment detection is not reducible to the family-pair space reported by HHsearch.

### 2.3 GPU benchmarking confirms numerical accuracy and database-scale feasibility

GPU benchmarking tested numerical agreement with CPU output, acceleration over a single CPU core, scaling on large Pfam subsets, and sparse output. GPU and CPU raw symmetric KL calculations agreed within the preset tolerances for all tested matrix elements: no element exceeded the 10^*−*4^ relative-error or 10^*−*5^ absolute-error limits, the maximum absolute error was 3.81 *×* 10^*−*6^, and the root mean square error was 5.07 *×* 10^*−*7^.

On larger GPU workloads, throughput increased with workload size. In the 10,000-pair benchmark containing 241.6 million window-pair comparisons, two RTX 3090 GPUs processed raw KL matrices in 153.47 s, corresponding to 1.57 *×* 10^6^ elements/s. For context, matched single-process CPU benchmarks fixed to one thread processed 20–500 family-pair workloads at 3.75–4.04 *×* 10^4^ elements/s. Thus, the large two-GPU benchmark operated at roughly forty-fold higher throughput than the measured single-thread CPU baseline, whereas smaller matched CPU/GPU benchmarks showed lower apparent speedups because fixed overheads are proportionally larger at small input size (Supplementary Methods Section S5 and Tables S7–S9). Linear extrapolation of the 10,000-pair two-GPU throughput to all unordered nonself Pfam family pairs gave an estimated 57.9 days.

The practical full-database workflow does not persist dense raw matrices. Instead, raw matrix batches are filtered immediately. In the 10,000-pair end-to-end benchmark, 241.6 million raw matrix elements were reduced to 30,862 first-stage candidates, 3,052 background-suppressed points, and 14 final paths. Thus, database-scale ProDive analysis is made practical by aggressive sparse filtering and path-level output, not by storing all raw window-pair values. Full benchmarking details, including memory measurements and dense-storage upper bounds, are provided in Supplementary Methods Section S5 and Tables S5–S12.

## 3 Discussion

Existing tools already excel at detecting family-level homology, domain architecture relationships, and structure-wide similarity.^5–8^ These tools answer different questions: whether whole domains, folds, or profile alignments are related, rather than systematically enumerating short local correspondences between protein-family fragments. ProDive addresses this problem using pHMMs as the current family representation; future representations of family-level local sequence distributions could be inserted into the same broader problem. ProDive is designed for this distinct fragment-scale detection problem: identifying local segment correspondences within the noisy all-window-pair landscape generated by two family models. The main technical result is that the closed-form divergence formulation can be embedded in a scalable workflow that produces pHMM segment correspondences across the full Pfam database and projects a structurally characterizable subset onto concrete chains.

The comparison with HHsearch clarifies that ProDive is not simply a boundary refinement of conventional profile–profile search. The central methodological difference is that HHsearch reports a path-based summary of profile–profile relatedness, whereas ProDive uses approximate full-profile probability scoring over short pHMM windows before linking local high-scoring windows into paths. Across both HHsearch thresholds, most ProDive paths occurred outside HHsearch-reported family-pair space, while most HHsearch-reported family pairs did not overlap a ProDive path. For family pairs reported by both methods, ProDive-defined boundaries isolated shorter compact cores, whereas HHsearch-defined boundaries usually described broader intervals. These observations establish non-overlapping value rather than a proven causal mechanism: HHsearch contributes broader profile-level relationship detection, whereas ProDive identifies local fragment pairs that are enriched for compact, lower-RMSD structural correspondence.

The present study also has limitations. First, the method depends on the quality and granularity of Pfam seed alignments and their derived pHMMs. Alternative family models or clustering strategies could change which local signals are detectable. Second, although window-length sensitivity was evaluated for the Pfam pHMM setting used here, the optimal window scale may depend on the family-model representation and on how diffusely local sequence preferences are distributed within a model. Extending this analysis to alternative model-building strategies, clustering schemes, or future representations of family-level local sequence distributions will be important. Other operating parameters, including path thresholds and seed-alignment-support cutoffs, were empirically chosen rather than learned from a goldstandard training set. Third, structural characterization is limited by the availability and accuracy of representative experimental structures or sufficiently confident AlphaFold predictions. As more experimental structures and improved predictions become available, additional ProDive segment correspondences can be projected onto concrete fragment pairs and structurally characterized without rerunning the detection stage. Fourth, our external benchmark is centered on HHsearch; broader comparisons against additional local or structure-based tools would be useful in future work. Finally, the current HHsearch comparison is observational. Future controlled analyses will be needed to disentangle the contributions of path-based profile-alignment optimization, distributional window scoring, threshold calibration, and downstream filtering to the observed overlap and boundary differences.

## 4 Conclusion

ProDive introduces an approximate full-probability pHMM-window divergence formulation for same-length profile segments and embeds it in a scalable Pfam-wide workflow. Applied to Pfam release 38.0, this framework identifies 318,289 high-confidence cross-family segment correspondences. Structural characterization shows that projected representative-chain fragments are enriched for compact low-RMSD local cores, and reciprocal comparison with HHsearch shows that ProDive captures a largely distinct fragment-scale signal rather than serving as a boundary refinement of domain-scale remote-homology search. These results establish ProDive as both a practical method and a large-scale resource for studying local sequence–structure conservation across protein families.

## 5 Methods

### 5.1 Closed-form window divergence

The main divergence formula is defined for equal-length pHMMs. In practice, a pHMM of length *n* is partitioned into *n−k* + 1 overlapping windows of length *k*, and a second pHMM of length *m* is partitioned analogously. The directional divergence *D*_∞_ is symmetrized by averaging the two directions:

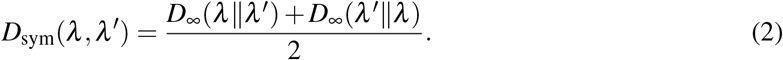

Because the divergence grows with window length, we normalize by the number of basic terms accumulated over the window and use

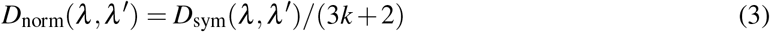

as the operational distance. All reported window-pair distances refer to *D*_norm_.

### 5.2 Candidate screening and background suppression

For two pHMMs A and B, ProDive evaluates all window-pair distances 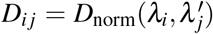 independently, which makes the calculation naturally parallelizable. A first-stage threshold *D*_*i j*_ *≤ ε* retains candidate pairs. In the production Pfam scan, we used *ε* = 0.1.

To suppress background-driven similarities, we computed how informative each pHMM match-state position was relative to the model background. For pHMM A, let *p*_*A,t*_(*a*) be the amino-acid emission probability at match-state position *t* and let *q*_*A*_(*a*) be the HH-suite `NULL` background distribution stored with that model. We defined the per-position background information as

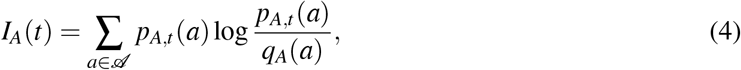

where *A* is the 20-amino-acid alphabet. For a window beginning at position *i*, the window information value was the average across its *k* matched positions,

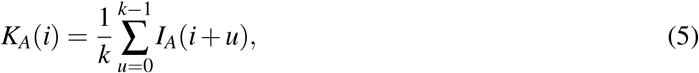

with *I*_*B*_(*t*) and *K*_*B*_( *j*) defined analogously for pHMM B. For a candidate point (*i, j*), these window information values were combined with the normalized divergence to form a background-suppressed score:

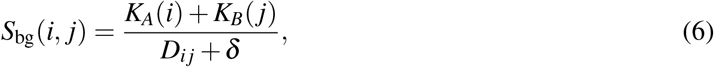

where *δ* is a numeric stabilizing constant. This favors pairs that are both locally informative and mutually similar, while down-weighting low-information background similarities.

### 5.3 Diagonal-path extraction and rescoring

Candidate points were linked into diagonal paths when both coordinates increased along the main diagonal direction and the step size was at most *k −* 1. Paths with fewer than five consecutive points were discarded in the production analysis. For each retained path, we recorded the Pfam family pair, the start and end pHMM state positions spanned on each family side after merging the overlapping windows in the path, the number of window-pair points supporting the path, and the mean background-suppressed point score 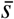 across those points. Retained paths were classified by assignment uniqueness: clean paths represented a one-to-one segment assignment, whereas ambiguous paths contained overlapping assignments and were excluded from downstream score-based analyses. Across the full Pfam scan, path extraction yielded 471,050 clean paths and 319,634 ambiguous paths.

For each pHMM state position, seed-alignment support was defined as the fraction of seed sequences carrying a residue at that position. For each path, *c*_main_ and *c*_sub_ denote the mean support across the matched segment on the main and sub family sides, respectively. These two values were combined into a weight *w* = (*c*_main_ + *c*_sub_)*/*2. We then rescored each path as

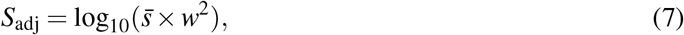

where 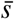 is the mean background-suppressed point score of the path. Paths were retained when *c*_main_ *≥* 0.85, *c*_sub_ *≥* 0.85, and *S*_adj_ *>* 1.2. Filtering-stage counts are summarized in Supplementary Methods Section S3 and Table S3.

### 5.4 Pfam-wide scan

All 25,545 families from Pfam release 38.0 were represented by seed-derived pHMMs. ^3^ The all-against-all scan was performed on sliding windows of length *k* = 6 using a custom GPU kernel to evaluate the independent window-pair distances. The full scan produced candidate matrices for all family pairs, from which diagonal paths were extracted and rescored as described above. A sensitivity analysis tested window lengths between 2 and 20. The production value *k* = 6 was the first setting that escaped the fragmented short-window regime while preserving much broader Pfam coverage than longer windows. Full results are provided in Supplementary Methods Section S2 and Tables S1–S2.

### 5.5 Performance benchmarking

Computation-cost analyses used the same family pairs, single-precision packed database, and raw symmetric KL output boundaries for CPU and GPU runs. CPU numeric libraries were fixed to one thread per process. Each reported runtime benchmark was repeated three times and summarized by the median unless otherwise stated. Throughput was computed in matrix elements per second, where one element corresponds to one window-pair comparison. For memory use, we recorded the peak host-memory footprint and the increase in allocated GPU memory relative to the pre-run baseline. End-to-end workflow measurements included KL calculation, two-stage threshold filtering, diagonal-path construction, and seed-alignment-support rescoring.

### 5.6 Structural characterization

Every retained path was projected from pHMM state positions to seed-alignment columns and then to representative protein sequences, producing one concrete fragment pair per segment correspondence.

Because insertions and deletions vary among sequences, the two realized fragment lengths could differ from each other and from the number of pHMM state positions in corresponding segments. A fragment pair entered structural characterization only when suitable representative structures were available for both fragments. Experimentally determined structures from the RCSB Protein Data Bank were preferred. ^19^ When no experimental structure was available, an AlphaFold model was used only when the fragment-average pLDDT *≥* 70 and endpoint PAE *≤* 10. ^20^ Fragment pairs were superposed with the Py-MOL `super` command, ^21^ and RMSD was calculated over aligned C*α* atoms. Reported aligned lengths and RMSD values therefore refer to this structurally characterized subset of representative-chain fragments, not directly to pHMM state positions. A coverage threshold of at least 0.8*L* aligned residues was required for a nominal pHMM segment length *L*. Structure availability, prediction-confidence, and coverage requirements were applied only after the ProDive segment correspondences and their projected fragment pairs had been determined; they affected inclusion in the structural analysis but not ProDive detection or filtering. Structural-characterization parameters and additional structural panels are provided in Supplementary Methods Section S4, Table S4, and Supplementary Figs. S1–S3.

### 5.7 Random controls

To assess whether observed cross-family local conservation could be explained by chance, we constructed length-stratified random controls. For each tested core length ℓ *∈* [8, 13], 25,000 random segment pairs were sampled independently from the same representative-structure pool used for the Pfam– Pfam structural characterization. Each random pair was assigned the same target length ℓ and processed through the same structure retrieval, quality filtering, coverage filtering, PyMOL superposition, and RMSD calculation pipeline as the ProDive fragment pairs. Real and random RMSD distributions were then compared separately for each ℓ using Welch’s *t*-test. ^22^

### 5.8 HHsearch benchmark

Pairwise Pfam–Pfam HHsearch was performed on the same family space used for the ProDive scan. ^8^ We evaluated both the default reporting threshold of 20% and a stricter 70% probability threshold used in prior fragment-reuse studies. ^11;13^ Pfam family pairs were grouped according to whether they were reported by ProDive only, HHsearch only, or both methods. For family pairs reported by both, structural distributions were characterized separately using ProDive-defined and HHsearch-defined pHMM segment boundaries, where HHsearch-defined boundaries refer to the intervals spanned by the reported profile–profile alignment path. For reciprocal overlap plots, a ProDive path and an HHsearch alignment in the same Pfam family pair were counted as shared when they satisfied a dual segment-coverage threshold of 0.2; the same rule was applied to ProDive- and HHsearch-denominator panels.

## Code availability

The ProDive source code is maintained in a private GitHub repository at https://github.com/putian74/prodive/tree/main/prodive.

## Data Availability Statement

Pfam seed alignments and family models are available from the Pfam database. ^3^ Structural characterization used publicly available entries from the RCSB Protein Data Bank ^19^ and AlphaFold structure predictions. ^20^ Derived ProDive path tables and processed structural-characterization summaries will be released with the code repository. Data derived in this study is available at https://doi.org/10.5281/zenodo.22009469.

## Acknowledgments

No funding was available for this study. No assistance from individuals who are not listed as authors was used.

