## supplemental info for "ProDive: detecting cross-family local conservation"

### Supplementary Notation

The table below summarizes the main symbols used in the derivation and in the window-based comparison framework. Unless otherwise stated, unprimed quantities refer to the first pHMM $\lambda$ and primed quantities to the second pHMM $\lambda'$.

| Symbol | Meaning | Used in |
|---|---|---|
| $\lambda, \lambda'$ | Parameter sets for the two equal-length pHMMs being compared. | Eqs. (S1), (S19) |
| $D_H(\lambda \| \lambda')$ | Heuristic KL-type divergence from pHMM $\lambda$ to pHMM $\lambda'$. | Eq. (S1) |
| $D_\infty(\lambda \| \lambda')$ | Asymptotic approximation of the heuristic divergence as the observation length tends to infinity. | Eq. (S19) |
| $S$ | Hidden-state set of a pHMM. | Eqs. (S1), (S3) |
| $\mathbf{V}$ | Observation alphabet (20 standard amino acids). | Eqs. (S1), (S3), (S20) |
| $N$ | Number of hidden states under the state convention used in the derivation; for a pHMM of length $n$, $N = 3n + 2$. | Eqs. (S2), (S8) |
| $n$ | pHMM length in state positions. | Eq. (S5) |
| $\boldsymbol{\pi}, \boldsymbol{\pi}'$ | Initial state-distribution vectors of $\lambda$ and $\lambda'$. | Eqs. (S1), (S11), (S20) |
| $\boldsymbol{\pi}_t$ | Initial probability vector after removing the absorbing End-state component. | Eqs. (S11), (S12), (S19) |
| $\mathbf{A}$ | Full transition matrix of the pHMM $\lambda$. | Eqs. (S1), (S4), (S8)–(S10) |
| $a_{ij}, a'_{ij}$ | Transition probabilities from hidden state $i$ to state $j$ in $\lambda$ and $\lambda'$. | Eqs. (S3), (S5) |
| $b_j(O), b'_j(O)$ | Emission probabilities of observation $O$ from state $j$ in $\lambda$ and $\lambda'$. | Eq. (S3) |
| $p_{A,t}(a)$ | Amino-acid emission probability for residue type $a$ at match-state position $t$ of pHMM A. | Main Methods |
| $q_A(a)$ | HH-suite `NULL` background probability for residue type $a$ in pHMM A. | Main Methods |
| $I_A(t)$ | KL background information of match-state position $t$, computed as $D_{\mathrm{KL}}(p_{A,t} \| q_A)$. | Main Methods |
| $K_A(i)$ | Mean background information across the length-$k$ window beginning at pHMM state position $i$ in pHMM A. | Main Methods |
| $\mathbf{W}$ | Column vector of state-specific divergence terms. | Eqs. (S1), (S2), (S9), (S11) |
| $W_i$ | State-specific divergence contribution associated with starting from state $i$. | Eqs. (S2), (S3) |
| $\mathbf{W}_t$ | Vector of state-specific divergence terms after removing the End-state component. | Eqs. (S11), (S12), (S19) |
| $\mathbf{I}$ | Identity matrix of appropriate dimension. | Eqs. (S1), (S4), (S14)–(S19) |
| $T$ | Observation-length parameter in the original matrix accumulation. | Eqs. (S1), (S4), (S8), (S9), (S14)–(S18) |
| $\Phi$ | Matrix accumulation term $\mathbf{I} + \mathbf{A} + \cdots + \mathbf{A}^{T-2}$. | Eq. (S4) |
| $\Phi_C$ | Matrix accumulation term $\mathbf{I} + \mathbf{C} + \cdots + \mathbf{C}^{T-2}$. | Eqs. (S14)–(S18) |

| Symbol | Meaning | Used in |
|---|---|---|
| $\mathbf{C}$ | Principal submatrix of the full transition matrix $\mathbf{A}$ after deleting the absorbing End-state row and column. | Eqs. (S6), (S10)–(S19) |
| $\mathbf{B}$ | Transition block from transient states to the absorbing End state in the profile-HMM block form. | Eqs. (S6), (S7), (S10) |
| $\rho(\mathbf{C})$ | Spectral radius of $\mathbf{C}$. | Eq. (S16) |
| $Q$ | Initial-observation correction term retained outside the closed-form matrix approximation. | Eqs. (S19), (S20) |
| $k$ | Sliding-window length used for local pHMM comparison. | Methods, Section S2 |
| $r$ | Minimum number of consecutive diagonal points required to define a path. | Methods, Section S2 |

### Supplementary Methods

#### Section S1. Derivation of the heuristic divergence approximation for equal-length profile HMMs

This section provides the detailed derivation of the equal-length profile-HMM comparison formula used in the main text. The derivation starts from the heuristic KL-type divergence proposed for HMMs by Mohammad and Tranter.[1] We use Markov-source KL-divergence-rate theory as a related reference point, but the profile-HMM case requires a separate derivation.[2] Because the transition matrix of a profile HMM contains an absorbing End state, the standard stationary-distribution simplifications used for recurrent Markov chains do not apply directly. We therefore re-derive the matrix-accumulation term by exploiting the topology of the profile-HMM transition matrix.

##### S1.1 Starting point

For two equal-length HMMs, the heuristic divergence can be written as

$$D_H(\lambda\|\lambda') = \boldsymbol{\pi}\left(\mathbf{I}+\mathbf{A}+\cdots+\mathbf{A}^{T-2}\right)\mathbf{W} + \sum_{o_1\in\mathbf{V}}\sum_{q_1}\pi_{q_1}b_{q_1}(o_1)\log\frac{\sum_{q_1}\pi_{q_1}b_{q_1}(o_1)}{\sum_{q_1}\pi'_{q_1}b'_{q_1}(o_1)}. \tag{S1}$$

Here $\mathbf{A}$ is the transition matrix, $\boldsymbol{\pi}$ is the initial state-distribution vector, and $\mathbf{W}$ collects state-specific divergence terms:

$$\mathbf{W} = \left(W_1,\ldots,W_N\right)^{\mathsf{T}}, \qquad W_i = W(q_2 \mid q_1 = i), \quad i = 1,\ldots,N. \tag{S2}$$

Under the notation used in this work, an equivalent normalized expression for $W_i$ is

$$W_i = \sum_{O\in\mathbf{V}} \left[ \sum_{j\in S} a_{ij} b_j(O) \log \frac{\sum_{j\in S} a_{ij} b_j(O)}{\sum_{j\in S} a'_{ij} b'_j(O)} \right]. \tag{S3}$$

For convenience we denote the matrix accumulation term by

$$\mathbf{\Phi} = \mathbf{I} + \mathbf{A} + \cdots + \mathbf{A}^{T-2}. \tag{S4}$$

For a profile HMM of length $n$, the state set contains Start, End, and match (M), insert (I), and delete (D) states associated with the profile positions, so the total number of hidden states is $N = 3n+2$. After ordering the states along the profile, the transition matrix $\mathbf{A}$ is upper triangular. Its diagonal elements are composed mainly of insert-state self-loops, zeros, and the End-state self-loop probability 1. For any insert state $I_k$,

$$0 \le a_{I_k I_k} < 1, \qquad \forall k \in \{1,2,\ldots,n\}. \tag{S5}$$

##### S1.2 Why the conventional Cesàro-limit route degenerates

One natural approach to simplifying $\mathbf{\Phi}$ is to use Cesàro averaging of matrix powers. In canonical block form, a Markov-chain transition matrix can be written as

$$\mathbf{P} = \begin{pmatrix} \mathbf{C} & \mathbf{B} \\ \mathbf{0} & \mathbf{\Gamma} \end{pmatrix}, \tag{S6}$$

with the associated Cesàro limit

$$\lim_{x\to\infty} \frac{1}{x} \sum_{i=1}^{x} \mathbf{P}^i = \begin{bmatrix} \mathbf{0} & (\mathbf{I}-\mathbf{C})^{-1}\mathbf{B}\mathbf{D}_{\text{proj}} \\ \mathbf{0} & \mathbf{D}_{\text{proj}} \end{bmatrix}. \tag{S7}$$

However, when this is applied directly to a profile HMM, End is the unique absorbing state and all other states are transient. The resulting limit for the full pHMM transition matrix produces a degenerate matrix term:

$$\lim_{T\to\infty} \frac{1}{T} \sum_{i=0}^{T-2} \mathbf{A}^i = \left[ \mathbf{0}_{N,N-1} \;\; \mathbf{1}_{N,1} \right]. \tag{S8}$$

Because the End state contributes no amino-acid emission term to **W**, the direct Cesàro-limit treatment collapses the leading matrix term in Eq. (S1) to zero:

$$\lim_{T\to\infty}\frac{1}{T}\boldsymbol{\pi}\left(\mathbf{I}+\mathbf{A}+\cdots+\mathbf{A}^{T-2}\right)\mathbf{W}=0. \tag{S9}$$

This degeneracy is the main reason a conventional recurrent-chain treatment is not suitable for the profile-HMM setting.

##### S1.3 Rewriting the matrix term using the transient-state block

The key observation is that the full matrix term can be rewritten using only the transient-state block of the profile-HMM transition matrix. Writing **A** in block form with transient-state submatrix **C** and the transient-to-End block **B** yields

$$\mathbf{A}^n=\begin{bmatrix}\mathbf{C}^n & \sum_{j=1}^{n}\mathbf{C}^{n-j}\mathbf{B}\\ \mathbf{0} & 1\end{bmatrix},\qquad n\geq 1. \tag{S10}$$

Multiplying by the initial probability vector and the emission-related vector gives

$$\boldsymbol{\pi}\mathbf{A}^n\mathbf{W}=\boldsymbol{\pi}_t\mathbf{C}^n\mathbf{W}_t,\qquad n\geq 1, \tag{S11}$$

where $\boldsymbol{\pi}_t$ and $\mathbf{W}_t$ are obtained by deleting the End-state component. Substituting Eq. (S11) into Eq. (S1) rewrites the heuristic divergence as

$$\begin{aligned}D_H(\lambda\|\lambda')=&\,\boldsymbol{\pi}_t\left(\mathbf{I}+\mathbf{C}+\cdots+\mathbf{C}^{T-2}\right)\mathbf{W}_t\\&+\sum_{o_1\in\mathbf{V}}\sum_{q_1}\pi_{q_1}b_{q_1}(o_1)\log\frac{\sum_{q_1}\pi_{q_1}b_{q_1}(o_1)}{\sum_{q_1}\pi'_{q_1}b'_{q_1}(o_1)}.\end{aligned} \tag{S12}$$

##### S1.4 Convergence of the transient-state series

Let

$$\Phi_C=\mathbf{I}+\mathbf{C}+\cdots+\mathbf{C}^{T-2}.$$

Since $\mathbf{C}$ is upper triangular, $\mathbf{I}-\mathbf{C}$ is also upper triangular and has strictly positive diagonal elements, so it is invertible:

$$\det(\mathbf{I}-\mathbf{C}) = \prod_i (1-C_{ii}) > 0. \tag{S13}$$

Using the matrix geometric-series identity,

$$\Phi_C(\mathbf{I}-\mathbf{C}) = \mathbf{I}-\mathbf{C}^{T-1}, \tag{S14}$$

and therefore

$$\Phi_C = (\mathbf{I}-\mathbf{C}^{T-1})(\mathbf{I}-\mathbf{C})^{-1}. \tag{S15}$$

Because the transient-state block contains only probabilities with absolute eigenvalues less than 1, its spectral radius satisfies

$$\rho(\mathbf{C}) < 1. \tag{S16}$$

Thus

$$\lim_{T\to\infty} \mathbf{C}^T = 0, \tag{S17}$$

and the transient-state accumulation converges to

$$\Phi_C \approx (\mathbf{I}-\mathbf{C})^{-1} \qquad \text{as } T\to\infty. \tag{S18}$$

##### S1.5 Final equal-length approximation

Substituting Eq. (S18) into Eq. (S12) gives the asymptotic approximation used in the main text:

$$D_\infty(\lambda\|\lambda') \approx \boldsymbol{\pi}_t(\mathbf{I}-\mathbf{C})^{-1}\mathbf{W}_t + Q, \tag{S19}$$

where

$$Q = \sum_{o_1\in\mathbf{V}}\sum_{q_1} \pi_{q_1} b_{q_1}(o_1) \log \frac{\sum_{q_1} \pi_{q_1} b_{q_1}(o_1)}{\sum_{q_1} \pi'_{q_1} b'_{q_1}(o_1)}. \tag{S20}$$

Equation (S19) is the theoretical basis of ProDive's equal-length comparison formula. Its importance lies in replacing the intractable finite-length matrix accumulation with the computable closed-form expression $\boldsymbol{\pi}_t(\mathbf{I}-\mathbf{C})^{-1}\mathbf{W}_t$.

#### Section S2. Sensitivity analysis supporting the choice of window length $k = 6$

We evaluated the production choice of sliding-window length $k = 6$ by repeating the window-pair and diagonal-path analysis on the same sampled set of 50,000 Pfam family pairs across $k = 2, 3, 4, 5, 6, 7, 8, 9, 10, 12, 14, 16, 18,$ The same family pairs were used for all values of $k$, so the differences reflect window length rather than different sampling backgrounds. Because some families have too few valid windows at larger $k$, the number of usable matrices was 50,000 for $k \leq 7$ and decreased modestly to 49,578 at $k = 20$.

Raw divergence scores increased approximately linearly with $k$, as expected for a quantity that accumulates over more HMM states. To compare across $k$, we therefore ranked candidates by low-distance percentiles after length normalization rather than by a single absolute raw threshold. The path-building criterion was adjusted to keep the minimum physical span close to 10 state positions. In this fixed-span setting, shorter windows require more consecutive diagonal points to form a path, whereas longer windows require fewer points (Table S1).

Table S1: Minimum diagonal-path point requirement used to keep the minimum physical span approximately comparable across window lengths.

| $k$ | 2 | 3 | 4 | 5 | 6 | 7 | 8 | 9 | 10 | 12 | 14 | 16 | 18 | 20 |
|---|---|---|---|---|---|---|---|---|---|---|---|---|---|---|
| Min path points | 9 | 8 | 7 | 6 | 5 | 4 | 3 | 3 | 3 | 3 | 3 | 3 | 3 | 3 |
| Approx. span | 10 | 10 | 10 | 10 | 10 | 10 | 10 | 11 | 12 | 14 | 16 | 18 | 20 | 22 |

Under a representative setting that retained the lowest 0.05% of length-normalized distances and used a second-stage background-suppression threshold of 10, short windows produced many retained points but very few continuous paths (Table S2). For example, $k = 2$ retained 695,714 candidate points but formed no paths, and $k = 3$ retained 413,217 points but formed only 36 paths. This indicates that very short windows are dominated by isolated or interrupted point-like similarities. At $k = 6$, the analysis first entered a practical regime: retained points decreased to 66,377, but path count rose to 1,211 and path conversion increased to 0.109 while still covering 15,945 Pfam families.

Longer windows showed higher path conversion rates, but this came with rapid loss of family-level coverage. At $k = 8$, covered Pfams dropped to 8,630; at $k = 10$, only 3,953 Pfams remained covered; and at $k = 12$, coverage fell to 1,637. The same trade-off was observed under stricter background filtering. Together, these results support $k = 6$ as a conservative working point rather than an absolute optimum. It is the first window length that substantially escapes the short-window regime while still preserving far broader Pfam coverage than longer windows.

Table S2: Representative window-length sensitivity results using the lowest 0.05% distance points and second-stage background-suppression threshold 10.

| $k$ | Min points | Retained points | Path count | Conversion | Noise fraction | Covered Pfams |
|---|---|---|---|---|---|---|
| 2 | 9 | 695,714 | 0 | 0.000 | 1.000 | 24,695 |
| 3 | 8 | 413,217 | 36 | 0.001 | 0.999 | 24,253 |
| 4 | 7 | 184,916 | 176 | 0.007 | 0.993 | 22,302 |
| 5 | 6 | 116,757 | 566 | 0.034 | 0.966 | 19,811 |
| 6 | 5 | 66,377 | 1,211 | 0.109 | 0.891 | 15,945 |
| 7 | 4 | 44,033 | 2,295 | 0.261 | 0.739 | 12,368 |
| 8 | 3 | 27,943 | 3,288 | 0.483 | 0.517 | 8,630 |
| 9 | 3 | 18,965 | 2,322 | 0.516 | 0.484 | 6,093 |
| 10 | 3 | 12,431 | 1,628 | 0.559 | 0.441 | 3,953 |
| 12 | 3 | 5,195 | 697 | 0.596 | 0.404 | 1,637 |

#### Section S3. Path filtering summary

Table S3 summarizes the main production filtering stages after diagonal-path extraction. These counts correspond to the clean one-to-one path set, the seed-alignment-support filter, and the final adjusted-score cutoff used to define the 318,289 high-confidence ProDive correspondences.

Table S3: Path counts across the main ProDive filtering stages in the Pfam-wide scan.

| **Stage** | **Count** | **Retained fraction** |
|---|---|---|
| Primary clean one-to-one paths | 471,050 | 1.000 |
| After completeness requirement | 452,080 | 0.960 |
| Final clean-path set after score cutoff | 318,289 | 0.676 |

#### Section S4. Structural-characterization parameters and additional panels

Table S4 summarizes the structure-availability, confidence, coverage, and random-control settings used for the structural characterization. The supplementary structural figures are placed here because they extend the same validation analyses reported in the main text.

Table S4: Structural-characterization pipeline parameters used in the ProDive methods manuscript.

| Parameter | Value | Description |
|---|---|---|
| AlphaFold pLDDT | $\geq 70$ | Fragment-average predicted confidence |
| AlphaFold PAE | $\leq 10$ | Endpoint predicted alignment error |
| Coverage requirement | $\geq 0.8L$ | Minimum aligned length as fraction of pHMM state-position number $L$ |
| Superposition method | PyMOL `super` | Structural superposition for RMSD calculation |
| Random control (Pfam–Pfam) | 25,000 pairs per length | Length-stratified sampling for $\ell \in [8, 13]$ |
| Statistical test | Welch's $t$-test | Comparison of real versus random RMSD distributions |

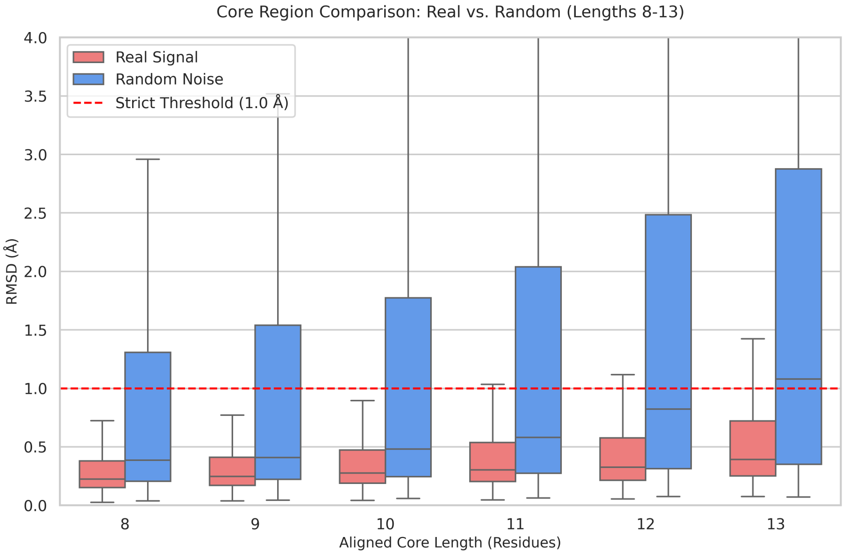


Figure S1: Length-stratified random control for the compact-core regime (8–13 residues). Boxplots compare RMSD distributions of structurally characterized representative-chain fragment pairs projected from ProDive segment correspondences in family pairs not reported by HHsearch against length-matched random fragment pairs for each aligned core length $\ell \in [8, 13]$.

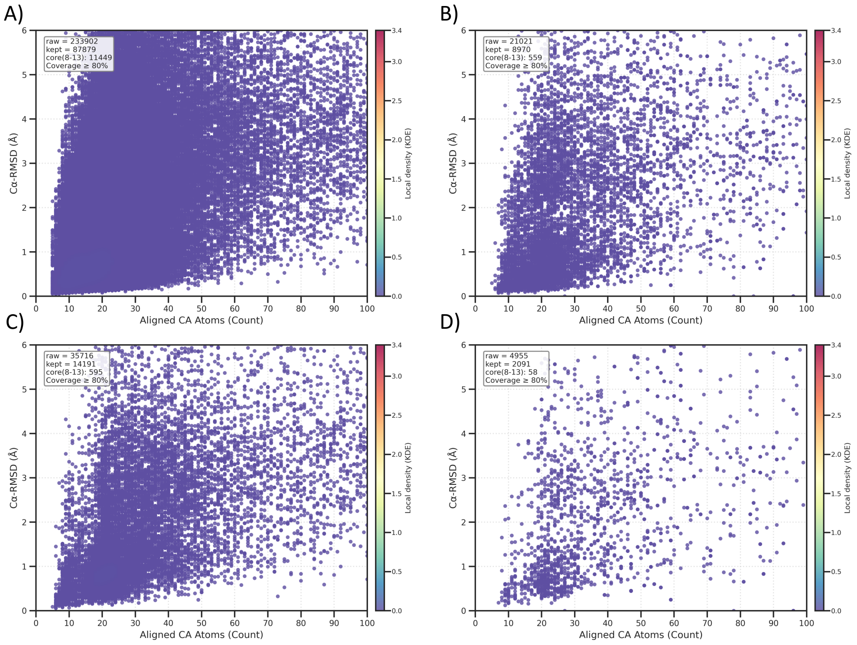


Figure S2: Full-range distributions of structurally characterized fragment pairs under HHsearch-defined pHMM segment boundaries. These plots use the same format as the main-text benchmark figure but extend the aligned-length axis to show the full span of longer fragments recovered using HHsearch segment boundaries.

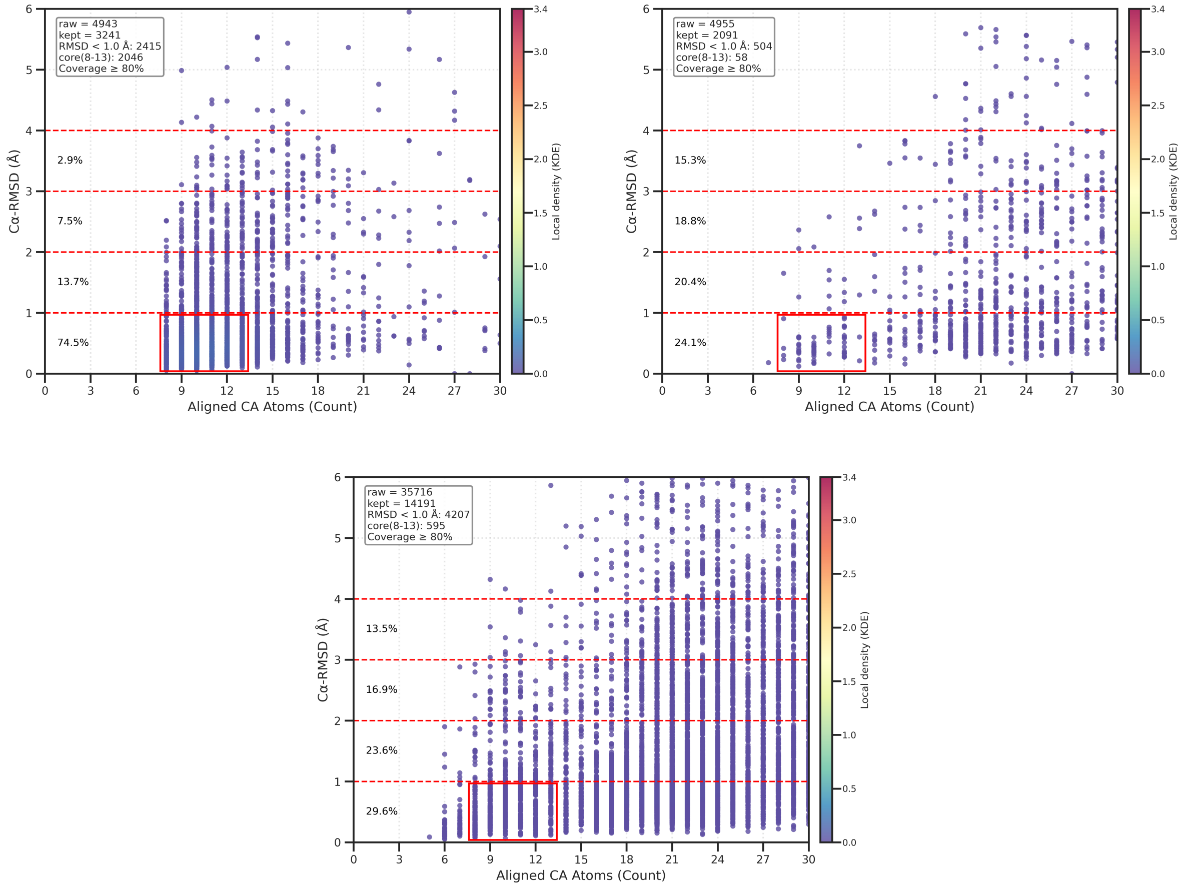


Figure S3: Additional structural-characterization panels at the stricter 70% HHsearch cutoff. All panels contain fragment pairs meeting the structural availability, prediction-confidence, and coverage requirements. (A) Representative-chain fragments from family pairs reported by both methods, extracted using ProDive-defined pHMM segment boundaries. (B) Representative-chain fragments from family pairs reported by both methods, extracted using HHsearch-defined pHMM segment boundaries. (C) Representative-chain fragments extracted from HHsearch-defined pHMM segments in family pairs without a ProDive path. These panels show that the contrast between compact fragments extracted using ProDive segment boundaries and broader fragments extracted using HHsearch segment boundaries persists under more stringent HHsearch thresholding.

#### Section S5. Computation cost and memory-footprint benchmark

Detailed benchmark results supporting the main-text performance summary are reported below, including hardware scale, numerical agreement, CPU/GPU runtime, memory footprint, sparse-output reduction, and full-database extrapolation. These tables also provide computation-cost context for the ProDive resource used by the companion biological-characterization manuscript.

Table S5: Hardware and database scale used for the ProDive performance benchmark.

| Item | Value |
|---|---:|
| CPU | Intel Xeon E5-2696 v3, 72 logical cores |
| Host memory | 62.76 GiB |
| GPU | $2 \times$ NVIDIA GeForce RTX 3090 |
| GPU memory | 24 GiB per card |
| Pfam families | 25,545 |
| Windows at $k = 6$ | 3,969,369 |
| Packed database size | 12.37 GB |
| Unordered non-self family pairs | 326,260,740 |
| All full-database matrix elements | 7,877,409,782,701 |

Table S6: Numerical consistency between CPU and GPU raw symmetric KL output.

| Metric | Value |
|---|---:|
| Family pairs | 5 |
| Compared matrix elements | 27,733 |
| Raw KL value range | 2.15–49.02 |
| Shape mismatches | 0 |
| NaN or infinite values | 0 |
| Relative-error tolerance | $10^{-4}$ |
| Absolute-error tolerance | $10^{-5}$ |
| Elements exceeding tolerance | 0 |
| Maximum absolute error | $3.81 \times 10^{-6}$ |
| Mean absolute error | $2.91 \times 10^{-7}$ |
| Root mean square error | $5.07 \times 10^{-7}$ |
| Pearson correlation | 0.9999999999999853 |

Table S7: Median raw-KL runtime, throughput, and acceleration in matched single-process CPU, one-GPU, and two-GPU comparisons.

| Pairs | Elements | CPU | 1 GPU | 2 GPUs | CPU throughput | 1-GPU speedup | 2-GPU speedup |
|---:|---:|---:|---:|---:|---:|---:|---:|
| 20 | 487,045 | 12.98 s | 0.93 s | 0.85 s | 37.54k/s | 14.02× | 15.33× |
| 50 | 1,091,934 | 29.05 s | 1.70 s | 1.31 s | 37.59k/s | 17.07× | 22.21× |
| 100 | 2,486,781 | 61.57 s | 2.85 s | 2.40 s | 40.39k/s | 21.64× | 25.65× |
| 200 | 5,009,275 | 128.33 s | 5.73 s | 4.04 s | 39.03k/s | 22.38× | 31.79× |
| 500 | 12,311,926 | 325.19 s | 13.78 s | 9.58 s | 37.86k/s | 23.59× | 33.95× |

Table S8: CPU multiprocessing benchmark on a fixed 500-pair workload (12,311,926 matrix elements).

| CPU processes | Median time | Throughput | Speedup | Efficiency | Host memory |
|---:|---:|---:|---:|---:|---:|
| 1 | 325.19 s | 37.86k/s | 1.00× | 100.0% | 0.86 GiB |
| 2 | 177.08 s | 69.53k/s | 1.84× | 91.8% | 1.52 GiB |
| 4 | 97.12 s | 126.77k/s | 3.35× | 83.7% | 1.85 GiB |
| 8 | 54.17 s | 227.29k/s | 6.00× | 75.0% | 2.51 GiB |
| 16 | 32.44 s | 379.51k/s | 10.02× | 62.7% | 3.84 GiB |

Table S9: GPU scaling, memory footprint, estimated energy use, and raw output.

| Pairs | Elements | GPUs | Median time | Throughput | Host memory | GPU-memory increment | Energy | Raw output |
|---:|---:|---:|---:|---:|---:|---:|---:|---:|
| 500 | 12,311,926 | 1 | 13.78 s | 893.16k/s | 0.75 GiB | 13.84 GiB | 0.77 Wh | 49.29 MB |
| 500 | 12,311,926 | 2 | 9.58 s | 1,285.23k/s | 0.83 GiB | 27.69 GiB | 0.91 Wh | 49.29 MB |
| 2,000 | 48,131,436 | 1 | 53.23 s | 904.14k/s | 2.53 GiB | 13.84 GiB | 2.90 Wh | 192.69 MB |
| 2,000 | 48,131,436 | 2 | 35.60 s | 1,351.84k/s | 2.62 GiB | 27.69 GiB | 3.40 Wh | 192.69 MB |
| 10,000 | 241,556,038 | 1 | 244.35 s | 988.57k/s | 9.28 GiB | 13.87 GiB | 12.88 Wh | 967.02 MB |
| 10,000 | 241,556,038 | 2 | 153.47 s | 1,573.99k/s | 9.36 GiB | 27.74 GiB | 13.49 Wh | 967.02 MB |

Table S10: End-to-end workflow cost for a fixed 10,000-pair input on two GPUs. The workflow includes KL calculation, two-stage thresholding, path construction, and completeness rescoring.

| Stage | Median time | Host memory | Output size |
|---|---:|---:|---:|
| KL calculation | 149.87 s | 9,589.4 MiB | 967.14 MB |
| Threshold filtering | 7.50 s | 250.6 MiB | 1.19 MB |
| Path construction | 17.99 s | 1,092.7 MiB | 4.57 KB |
| Completeness rescoring | 1.95 s | 188.4 MiB | 163.19 KB |
| Complete workflow | 177.58 s | — | — |

Table S11: Data reduction during the 10,000-pair end-to-end benchmark.

| Processing node | Count | Fraction of raw elements | Reduction |
|---|---:|---:|---:|
| Raw matrix elements | 241,556,038 | 100% | 1× |
| First threshold retained | 30,862 | 0.012776% | 7,826.97× |
| Second threshold retained | 3,052 | 0.001263% | 79,146.80× |
| Final paths after rescoring | 14 | — | — |

Table S12: Linear extrapolation from the 10,000-pair raw-KL benchmark to the full unordered non-self Pfam family-pair space. The dense raw float32 payload is a hypothetical storage upper bound if all matrix elements were retained; the operational workflow instead persists sparse filtered candidates and paths.

| GPUs | Measured throughput | Extrapolated time | Dense raw float32 payload |
|---:|---:|---:|---:|
| 1 | 988,568.98 elements/s | 92.23 days | 31.51 TB |
| 2 | 1,573,991.43 elements/s | 57.93 days | 31.51 TB |
